# Genome-wide abundance of 5-hydroxymethylcytosine in breast tissue reveals unique function in dynamic gene regulation and carcinogenesis

**DOI:** 10.1101/339069

**Authors:** Owen M. Wilkins, Kevin C. Johnson, E. Andres Houseman, Jessica E. King, Carmen J. Marsit, Brock C. Christensen

## Abstract

5-hydroxymethylcytosine (5hmC) is generated by oxidation of 5-methylcytosine (5mC), however little is understood regarding the distribution and functions of 5hmC in mammalian cells. We determined the genome-wide distribution of 5hmC and 5mC in normal breast tissue from disease-free women. Although less abundant than 5mC, 5hmC is differentially distributed, and consistently enriched among breast-specific enhancers and transcriptionally active chromatin. In contrast, regulatory regions associated with transcriptional inactivity were relatively depleted of 5hmC. Gene regions containing abundant 5hmC were significantly associated with lactate oxidation, immune cell function, and prolactin signaling pathways. In independent data sets, normal breast tissue 5hmC was significantly enriched among CpG loci demonstrated to have altered methylation in pre-invasive breast cancer and invasive breast tumors. Our findings provide a genome-wide map of nucleotide-level 5hmC in normal breast tissue and demonstrate that 5hmC is positioned to contribute to gene regulatory functions which protect against carcinogenesis.

## Introduction

5-methylcytosine (5mC) is the predominant cytosine modification of the mammalian genome, and a major coordinator of tissue-speific gene expression programs^1^. Recent studies demonstrate 5mC can be oxidized to several distinct chemical entities^2, 3, 4^, which have been observed to be abundant and stable under particular biological contexts^5^. There is an emerging appreciation that these oxidized forms of 5mC contribute actively to gene regulation and chromatin structure, however our knowledge of tissue specific distribution and functions of oxidized 5mC is limited.

5-hydroxymethylcytosine (5hmC), the most abundant form of oxidized 5mC^4^, is the first intermediate produced through oxidation of 5mC by the ten-eleven translocation (TET) family of enzymes^2, 3^. Enrichment of 5hmC among enhancers, DNase hypersensitivity sites, transcription factor binding sites, and sense-strand DNA suggest a positive association with transcription^6, 7, 8, 9, 10^. Proteins capable of binding 5hmC have been identified and suggest enrichment for proteins involved in transcription and chromatin regulation^11, 12, 13^. Co-localization of 5hmC at sites of DNA damage has promoted discussion about its potential functions in the coordination of DNA repair and genomic stability^14, 15^. Contrasting with that of 5mC, distribution and abundance of 5hmC varies greatly across human tissues. Brain and breast tissue possess the most abundant 5hmC among human tissues^16^. Approximately four times greater levels of 5hmC has been observed in brain relative to breast tissue^16^, and brain has therefore been the focus of early studies characterizing the distribution and potential functions of 5hmC in human tissues^2, 17, 18, 19^. However, less is known regarding the genomic distribution and functions of 5hmC in breast tissue.

During development, DNA methylation is responsible for coordination of tissue polarity and cellular differentiation of the mammary epithelium^20^. Dysregulation of DNA methylation is an early event in breast carcinogenesis, and methylation changes between normal and malignant breast tissue are widespread^21, 22^. Reduction of total 5hmC measured using global approaches has been observed in breast cancer, as well as most other tumor types^23, 24^. Indeed, decreased levels of 5hmC have been reported as a poor prognostic factor in breast cancer^25^. Consistent with these data, numerous mechanisms that abrogate *TET* expression and function have been observed in human cancer^26, 27, 28^. These observations strongly suggest tumor suppressive functions for active DNA demethylation and 5hmC and underscore the need to more thoroughly understand epigenetic regulation through study of oxidized 5mC.

Here, we utilize paired bisulfite (BS) and oxidative-bisulfite (oxBS) DNA modification followed by assessment using Infinium DNA methylation arrays to produce a genome-wide map of 5mC and 5hmC distribution in normal breast tissue. While 5hmC is less abundant relative to 5mC throughout much of the breast genome, we identified specific regions containing abundant 5hmC. Furthermore, we demonstrate that the distribution of high 5hmC is dependent on genomic-context, and enriched in gene regulatory regions associated with open chromatin and active transcription. Finally, we demonstrate 5hmC enrichment among regulatory regions in transformed and malignant breast cancer cell lines, and observed 5hmC is enriched among CpGs differentially methylated between normal and tumor tissue.

## Results

### 5hmC is abundant and uniquely distributed in normal breast tissue

To assess the potential functional importance of 5hmC in normal breast tissue, we measured genome-wide 5hmC and 5mC in normal breast tissue samples from 18 healthy female volunteers. All tissue samples were from female subjects across a wide range of ages and BMI values. Study subjects were aged between 13 and 81 years (median = 54.5 years), weighed between 41-204 kg (median = 68 kg) and had BMI values between 15-63 (median = 25). To obtain total and single-nucleotide resolution estimates for 5hmC and 5mC we applied the OxyBS algorithm^29^ to data from paired BS and oxBS treated DNA measured with 450K arrays. Altered expression of enzymes that process cytosine modifications on DNA (such as *TET1, TET2, DNMT3A* and *DNMT3B*) could potentially explain the observed variation in total 5hmC across samples, therefore we tested for correlation between total 5hmC and expression of enzymes that process cytosine modification. Generally, there was no correlative evidence between total 5hmC and gene expression of cytosine modifying enzymes, however we did observe modest correlation between total 5hmC and *DNMT3A* expression (**r**^2^=0.52, *P*=0.03, **Supplementary Table 1**). Supporting the hypothesis that 5hmC may have function independent of its role in regeneration of naïve cytosine, we tested the association between total 5hmC (that is, the sum of 5hmC β-values across all CpGs in a sample divided by the total number of CpGs profiled) and 5mC content across all samples and no evidence of a significant relationship was observed (linear regression *P* = 0.11, **Supplementary Figure 1**). Variation in cell-type composition of heterogeneous tissues is known to be a potential confounder in epigenomic studies^30^, and could impact 5hmC levels. To address this issue, we estimated putative cell types and their proportions using a reference-free cell-type deconvolution-based approach (RefFreeEWAS)^31^, which suggested two major cell populations with varying proportions across subjects in these samples (**Supplementary Figure 2**). However, no relationship was observed between total 5hmC content and cellular proportions (**Supplementary Figure 2**).

We next sought to evaluate the genome-wide distribution of 5hmC at the nucleotide level. Cumulative density plots of mean 5hmC and 5mC at each measured CpG across all samples revealed that the majority of CpGs had very low levels of 5hmC and that mean 5mC levels across all CpGs had the expected bimodal distribution (**Figure 1A**). Most CpGs also showed a negative correlation between 5hmC and 5mC levels, with only approximately 10% of all CpGs demonstrating a positive correlation (**Figure 1B**). We observed the well-documented characteristic dependency of CpG island context on 5mC abundance, with relatively lower proportions of 5mC situated within CpG island and shore regions compared to ocean and shelf regions (**Figure 1C**). Despite substantially lower abundance of 5hmC, a similar pattern of abundance was observed across the CpG island context strata to that of 5mC, with the lowest levels of 5hmC present within CpG island and CpG island shores (**Figure 1D**). Functions of DNA methylation are known to vary by genomic region, therefore we posited that 5hmC may be differentially distributed among CpG islands, shores, shelves, and oceans in breast tissue. The highest levels of both 5hmC and 5mC were observed within CpG island shelf regions (**Figures 1C & 1D**). Similar levels of 5hmC was seen across CpG oceans, shores and shelves while shores exhibited greater levels of 5mC relative to oceans and shelves (**Figure 1D**).

**Figure 1.**
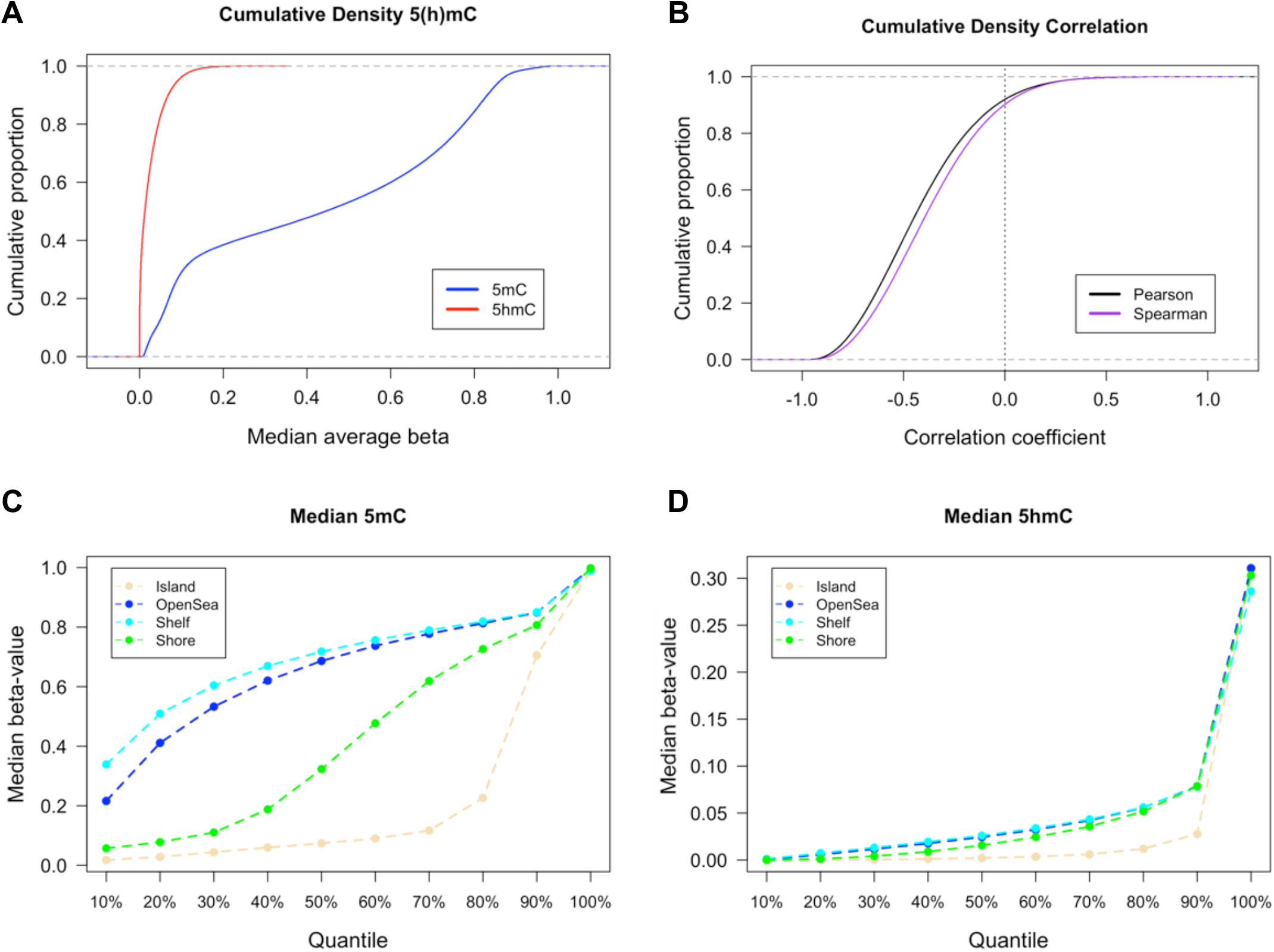
Abundance and genomic distribution of 5hmC in breast tissue. (**A**) Cumulative density distributions of median 5hmC and 5mC methylation beta-values breast tissue (*n* = 18). (**B**) Cumulative density distributions of Pearson and Spearman correlation coefficients as calculated for the relationship between 5hmC and 5mC beta-values at each CpG loci across all breast tissues (*n* = 18). Percentiles of median 5mC (**C**) and 5hmC (**D**) beta-values as calculated across all breast tissues and stratified by CpG island context (islands, shores, shelves, and oceans).

### Genomic enrichment of 5hmC in normal breast tissue

To better understand the potential functional relevance of 5hmC in breast tissue, we sought to identify CpG loci with the highest consistent abundance across samples. Large numbers of CpGs had median 5hmC beta-values near 0, though 66,341 CpG sites had median 5hmC levels above 5%, 14,733 CpG sites had median 5hmC levels above 10%, and 2,881 CpG sites had median 5hmC levels above 15% (**Figure 2A**). Aiming to distinguish functional 5hmC signal from background noise due to the generally sparse nature of 5hmC abundance, we selected 3876 CpGs with the highest 1% median 5hmC values, as calculated across all samples (**Figure 2B**). These CpGs had a range of median 5hmC beta-values from 14% to 31% and were defined as the ‘high 5hmC CpGs’ utilized in subsequent analyses. We utilized the quantile distribution of 5hmC beta-values to measure the consistency of each high 5hmC CpG across all tested breast tissues. 84.6% of the high 5hmC CpGs (3281/3876) presented with a 5hmC beta value of >0.15 (25^th^ percentile of 5hmC beta-value distribution) in at least half of the 18 samples, demonstrating these CpG loci were consistently elevated across samples. In contrast, 12.9% (500/3876) of the high 5hmC CpGs were among the top 1% most variable, based on standard deviation, suggesting more inter-subject variability. Basic genomic annotation data, as well as CpG-specific median and standard deviation are provided for the complete high 5hmC CpG set in **Supplementary Data 1**. To assess the relationship between 5hmC and 5mC abundance at a CpG-specific level, we compared 5hmC and 5mC beta-values at all high 5hmC CpG sites across all subjects. Generally, the greatest levels of 5hmC were observed for CpGs with intermediate to low levels of 5mC (**Figure 2C**), suggesting 5hmC may function in a manner distinct from 5mC. The majority of high 5hmC CpGs (3342/3876, 86.2%) were located within promoters, exons or introns), with the remaining CpGs located in intergenic regions (**Supplementary Data 2**). *SEPT7*, a key cytoskeletal gene, had a total of 9 high 5hmC CpGs located either within or proximal to the gene (**Supplementary Data 2**). In addition, there were 44 genes with at least 5 high 5hmC CpG located within or proximal to them. An overwhelming proportion of these the high 5hmC CpGs associated with these genes were located within intronic regions (**Supplementary Data 2**), consistent with previous findings that 5hmC is highly enriched at gene bodies^9, 32^. In addition, many of the genes containing at least 5 high 5hmC CpGs were located within transcriptional (co)activators or chromatin modifying genes, including *NCOR2, ARID1B, DNMT3A, FOXO3*, and *FOXP1*, among others (**Supplementary Data 2**). Given the positive correlation between 5hmC and gene expression that has been observed across many studies, presence of 5hmC at tumor suppressor genes (TSGs) may be an important regulatory mechanism is ensuring their expression to prevent carcinogenesis. Indeed, several genes with several high 5hmC CpGs located within their gene bodies are considered tumor suppressor genes (TSGs) in breast tissue, including *MBNL1, ARID1B, DNMT3A*, and *FOXO3*. Unsupervised clustering analyses using the high 5hmC loci did not clearly distinguish subjects by age, BMI or cell type proportion (**Supplementary Figure 3**).

**Figure 2.**
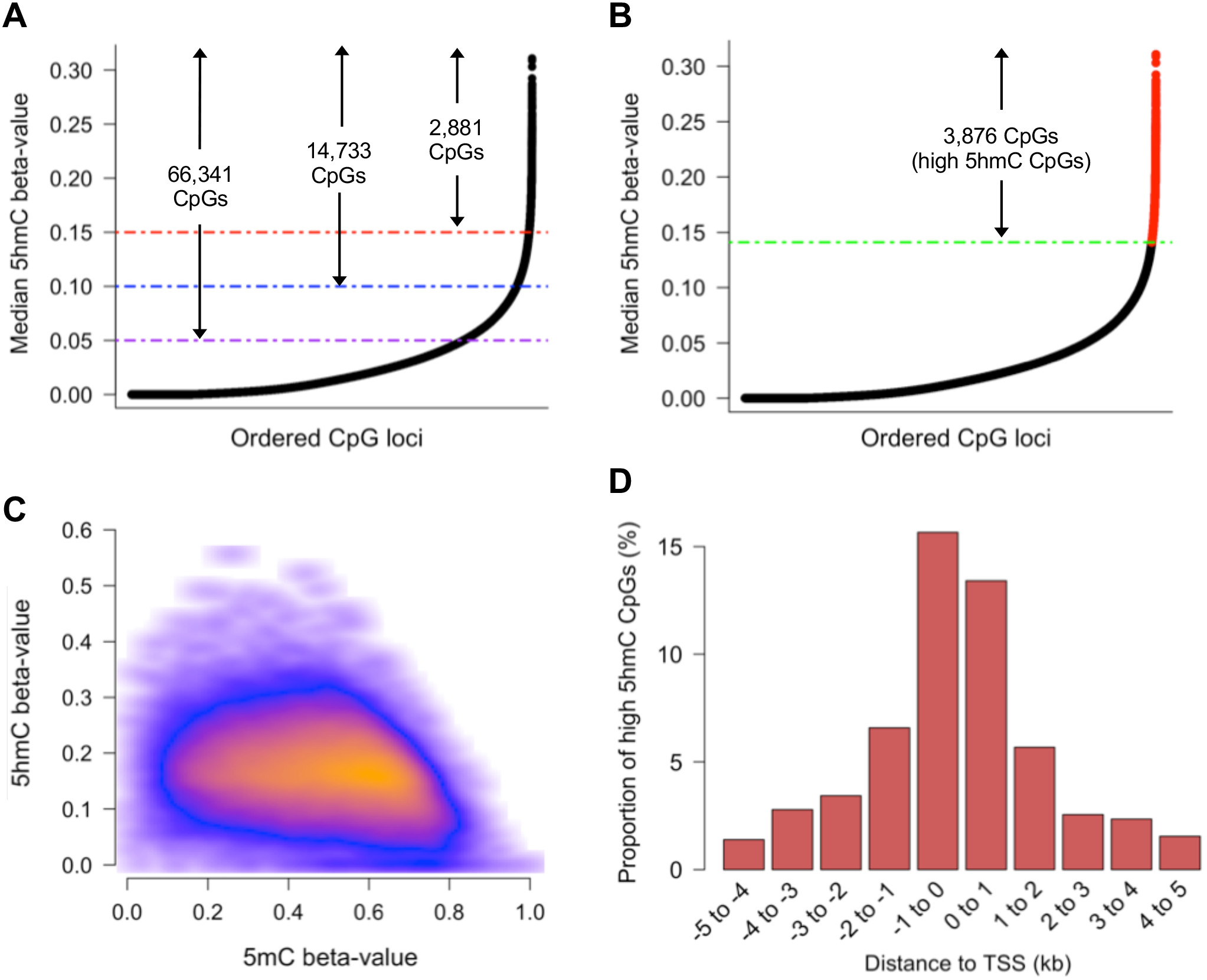
5hmC is uniquely distributed in breast tissue. (**A**) Rank ordered distribution of CpG-specific median 5hmC as calculated across 18 breast tissues. Purple, blue, and red lines, in conjunction with arrows and labels, indicate number of CpG loci with at least a minimum beta-value of 0.05, 0.10, and 0.15, respectively. (**B**) Rank ordered distribution of CpG-specific median 5hmC as calculated across 18 breast tissues, with the highest 1% mean 5hmC values (the ‘high 5hmC CpG sites’) across all samples denoted in red. Green line denotes mean 5hmC value of the 3876^th^ rank ordered high 5hmC CpG. (**C**) Scatter density plot of 5hmC beta-value vs 5mC beta-value for all high 5hmC CpGs. Each high 5hmC CpG is plotted once for each sample. Regions of orange and red indicate higher density of CpGs, whereas darker (black) regions indicate sparsity. (**D**) High 5hmC CpG site distribution relative to nearest canonical transcriptional start site (TSS). Vertical axis indicates percentage of high 5hmC within each distance grouping denoted on horizontal axis. Distance groupings provided for regions upstream and downstream on canonical TSSs. kb; kilo bases.

Previous studies in both non-diseased and diseased tissue have suggested an enrichment for 5hmC amongst regions involved in transcriptional regulation and gene bodies^6, 9, 32^. Amongst the high 5hmC CpGs, similar proportions were observed up and downstream (51.7% and 48.3%, respectively) of the nearest canonical transcriptional start site (TSS) (**Supplementary Figure 4**), with the majority situated within at most 1kb of the TSS (**Figure 2D**). To explore the potential functions of 5hmC, we tested enrichment of 5hmC among genomic features (promoters, exons, introns, and intergenic regions) and breast specific DNA regulatory regions using the Cochran-Mantel-Haenzel test to adjust for CpG context due to the varying proportion of 5hmC observed across CpG island, shores, shelves, and oceans. Consistent with previous findings^33, 34, 35^, we observed a substantial enrichment for intronic 5hmC, yet a dearth of exonic and intergenic 5hmC. To test the enrichment of 5hmC among breast specific regulatory regions, we utilized publicly available ChIP-seq data for a variety of histone modifications in normal breast myoepithelial cells and human mammary epithelial cells (HMECs). 5hmC was enriched in regions of active chromatin, marked by H3K4me3 in breast myoepithelial cells (**Figure 3**). Depletion of 5hmC was observed among H3K9ac-marked DNA, indicative if active promoters, and H3K9me3 and H3K27me3-marked DNA, indicative of transcriptionally inactive chromatin (**Figure 3**). In HMECs, a substantial enrichment of 5hmC was observed at poised and active enhancer regions, marked by H3K4me1 (OR; 3.2, 95% CI; 3.0-3.4, **Figure 3**). A smaller yet sizable enrichment of 5hmC at H3K27ac-marked regions (OR; 1.58, 95% CI; 1.47-1.71, Figure 3), which distinguishes active enhancers from those that are inactive or poised, suggested that more breast-specific enhancer 5hmC may be located within poised enhancers. Furthermore, regions of markers of transcriptionally active regions (H3K4me3, H3K36me3, H3K79me2, H3K20me1) also displayed positive enrichment for 5hmC (**Figure 3**). Conversely, 5hmC was depleted among repressed and transcriptionally inactive chromatin regions, marked by H3K9me3 and H3K27me3. DNase hypersensitivity sites and H2A.Z-marked regions did not show a significant enrichment for 5hmC in HMECs (**Figure 3**). Together, these results demonstrate breast-specific 5hmC is strongly enriched among gene regulatory regions associated with transcription, while severely depleted among regions associated with transcriptional repression.

**Figure 3.**
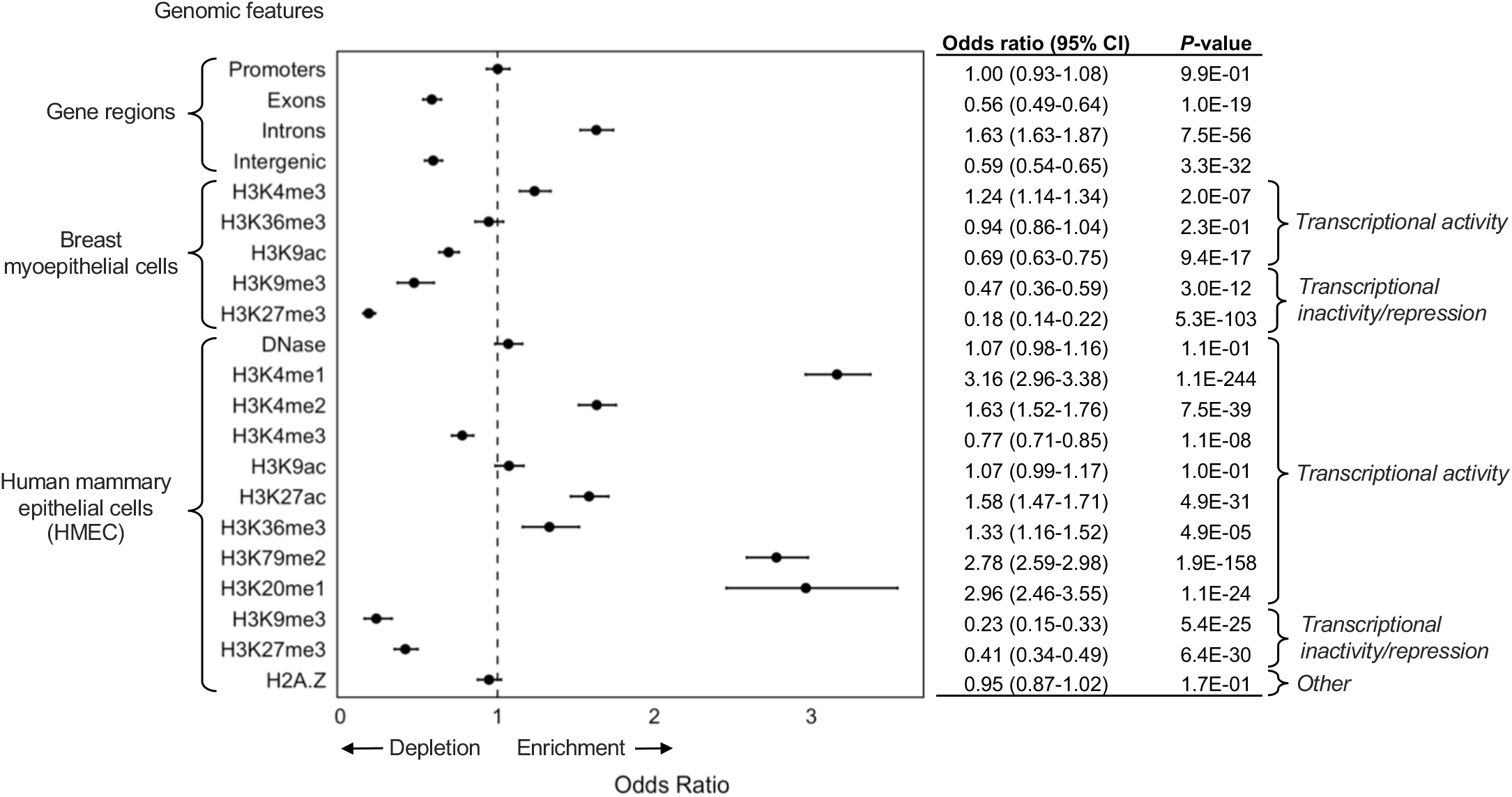
5hmC is enriched at regions associated with transcriptional activity. Forest plot of Cochran-Mantel-Haenszel odds ratios and 95% confidence intervals (95% CI) from testing enrichment of high 5hmC CpGs among indicated genomic features, against 450K background set. Numerical values for odds ratios and *P*-values are provided in the adjacent table.

### Gene set enrichment of 5hmC

While 5hmC is enriched among regulatory regions in breast tissue, it is plausible that a higher order structure exists for its distribution such that 5hmC is involved in regulation of specific gene sets. To investigate potential enrichment of 5hmC among specific gene regulatory programs, genomic coordinates of the high 5hmC CpGs were tested for gene set enrichment using the Genomic Regions Enrichment of Annotations Tool (GREAT). CpG loci measured on the 450K array were used as the background test set in these analyses. The ten most enriched gene sets with FDR Q-values < 0.05 from each of the; i) biological process, ii) molecular function, and iii) cellular process gene ontology analyses are presented in **Table 1**. Results for all gene sets significant at FDR < 0.05 are provided in **Supplementary Data 3**. The most enriched gene set in the biological process analysis was for lactate oxidation (fold enrichment; 28.3, hypergeometric FDR Q-value; 0.03, **Table 1**). Among the most enriched gene sets were also those relating to production of interleukins, function and activity of various immune cell types, and several biosynthetic processes including D-serine synthesis (**Table 1**). Interestingly, enrichment of the prolactin signaling pathway gene sets suggests potential involvement of 5hmC in regulating breast milk production (**Table 1**). Enrichment of molecular function gene sets relating to interleukin binding and D-serine biosynthesis activity further supported the results observed in the analyses of biological process gene sets. Enrichment was also observed for several molecular function gene sets related to binding and activity of TGF-beta (**Supplementary Data 4**), supporting the observation that five of the high 5hmC CpGs were located within the gene body of *TGFBR2* (**Supplementary Data 4**). Consistent with recent findings that 5hmC may be involved in the regulation genome stability, the most strongly enriched gene set in the cellular process ontology analysis was related to the replication fork protection complex (fold enrichment; 28.3, hypergeometric FDR Q-value; 0.02, **Table 1**). In addition, several of the enriched cellular process gene sets related to cell tip and membrane regulation (**Table 1, Supplementary Data 5**).

**Table 1.**
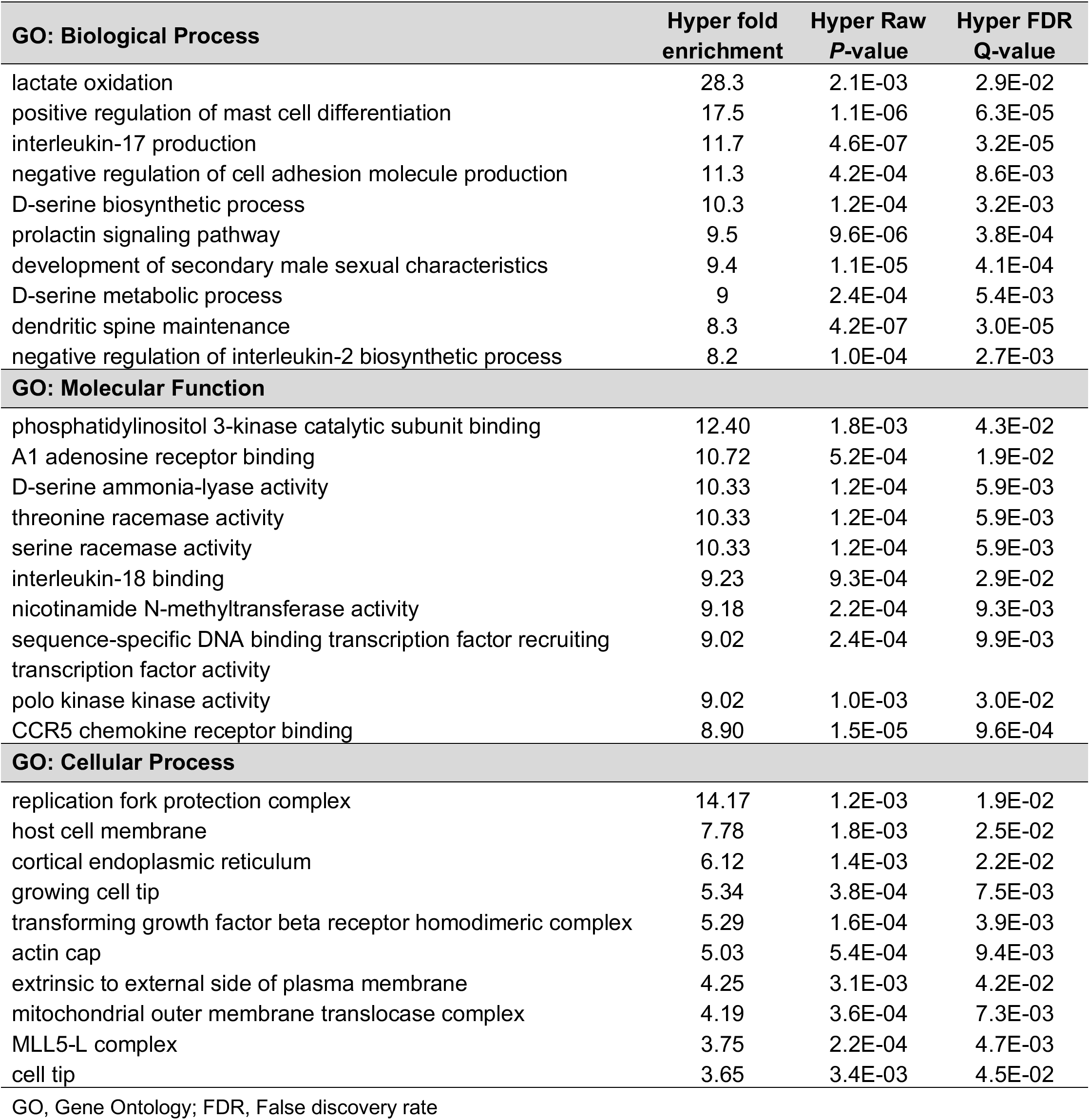
Genomic Regions Enrichment of Annotations Tool (GREAT) analysis of genomic regions containing high breast 5hmC CpGs

### Relationship between 5hmC and gene expression in breast tissue

Numerous studies have observed enrichment for 5hmC among regions of transcriptionally active chromatin^6, 9, 32^, and it is thought that 5hmC contributes to the positive regulation of gene expression in these regions^5^. To test the relationship between 5hmC abundance and gene expression, we measured transcript levels of four genes with known functions in breast tissue and multiple high 5hmC CpG loci. Generally, 5hmC was positively correlated with expression across the measured transcripts (**Figure 4A, Supplementary Data 7**), while 5mC showed mostly negative correlations with gene expression (**Figure 4B, Supplementary Data 7**). Among the most statistically significant correlations were those of cg23267550 and cg01915609 with *RAB32* expression for both 5hmC and 5mC (**Figures 4D & 4E**). The remaining four high 5hmC CpGs associated with *RAB32* demonstrated positive correlations between 5hmC and gene expression, while exhibiting negative correlations between 5mC and gene abundance, but did not reach statistical significance (**Figures 4A & B**). All six of the high 5hmC CpGs associated with *RAB32* were located in the within 1500bp of the TSS (TSS1500), at CpG island shore regions (**Figure 4C**). Across the transcripts measured for breast cancer tumor suppressor *RASSF1A*, there was evidence for positive correlations between 5hmC and expression for three CpG-expression relationships, while almost all of the tested CpGs showed negative correlations between expression and 5mC (**Supplementary Data 7**). Although several studies have suggested roles for 5hmC in regulation of alternative splicing, we did not observe dramatic modification of 5hmC-gene expression correlations by transcript-specific expression of RASSF1 and DNMT3A, suggesting 5hmC may not regulate mRNA splicing at these genes in breast tissue (**Supplementary Data 8**).

**Figure 4.**
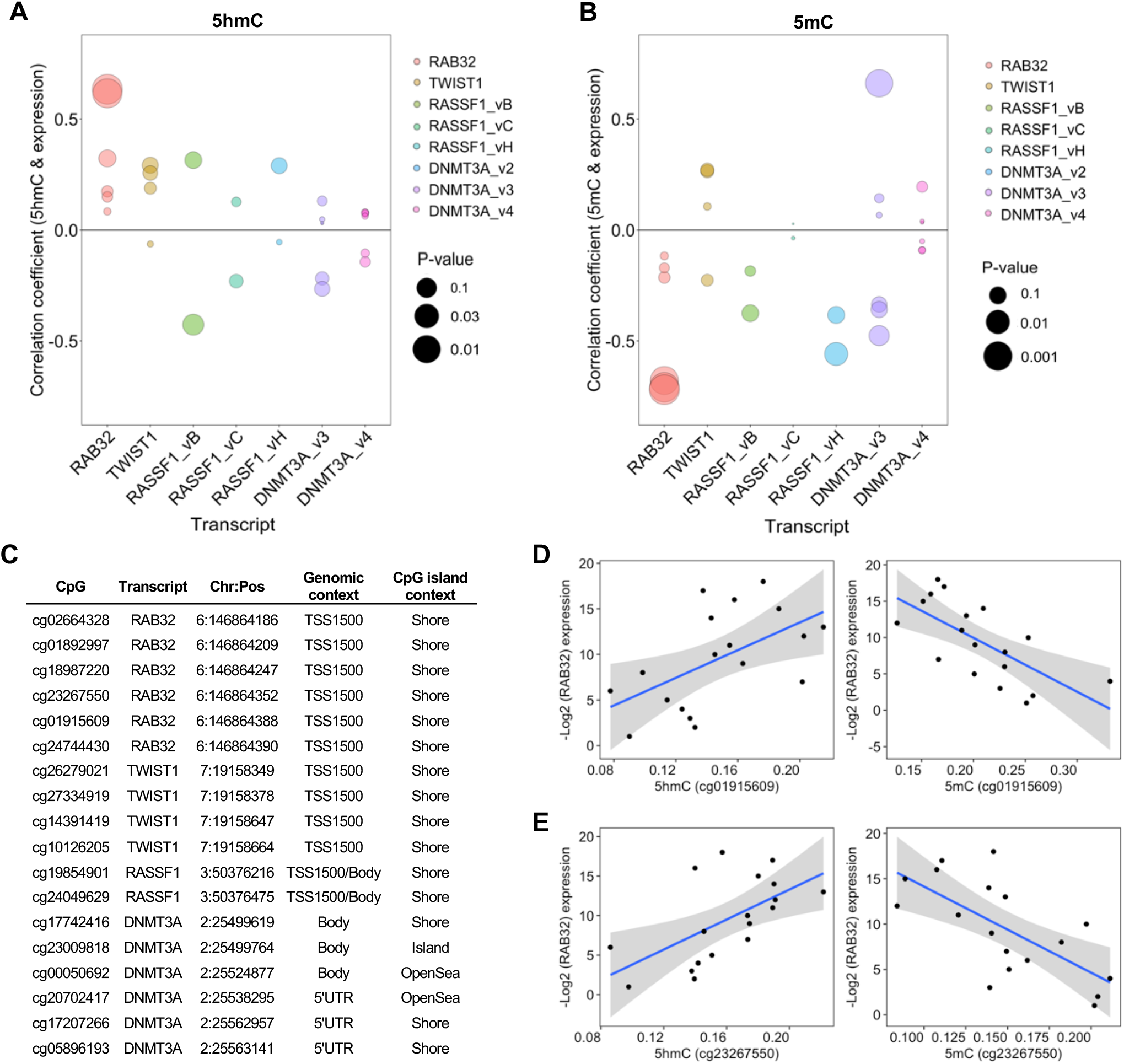
5hmC is positively associated with gene expression. Spearman correlation coefficients of relationship between 5hmC (**A**) and 5mC (**B**) abundance and gene expression at high 5hmC CpGs associated with candidate genes of interest. Colors denote methylation-expression relationship of CpGs associated with the genes denoted on x-axis. Bubble size corresponds to statistical significance (-Log_10_ *P*-value) associated with correlation coefficient. (**C**) Genomic annotation for the 18 high 5hmC CpGs tested for correlations between methylation and gene expression. (**D & E**) Scatter plots of subject-specific 5hmC or 5mC beta-values for cg01915609 (d) and cg23267550 (e) against Log_2_ expression values for *RAB32*. Regression line indicated in blue with 95% confidence bands in gray.

### Normal breast tissue 5hmC is enriched at regulatory regions in transformed and malignant breast cancer cell lines

Epigenetic deregulation is an early event in carcinogenesis. Substantial deviation from DNA methylation patterns in normal breast tissue have been observed in premalignant lesions as well as advanced disease. Although 5hmC tends to be depleted in proliferating cells, recent studies have suggested 5hmC may actively contribute to cancer pathogenesis. To determine whether 5hmC may contribute to breast carcinogenesis, we tested the normal breast tissue ‘high 5hmC CpGs’ for enrichment among known regulatory regions in variant human mammary epithelial cells (vHMECs). vHMECs are proliferative clones of regular HMECs that invariably result during cell culture, and are considered a model of premalignant breast cancer. Strong enrichment for the normal breast tissue high 5hmC CpGs was observed among regions marked by H3K4me1, associated with distal enhancer regions, as well as H3K4me36, associated with active transcription (**Figure 5A**). The strongest depletion of the normal breast tissue high 5hmC CpGs in vHMECs was observed among regions marked by H3K9me3 and H3K27me3, associated with transcriptional repression and inactive chromatin. Depletion was also observed at H3K4me3-marked regions, generally associated with gene activation. Regions of high normal breast tissue 5hmC were not enriched among DNase hypersensitivity sites in vHMECs.

**Figure 5.**
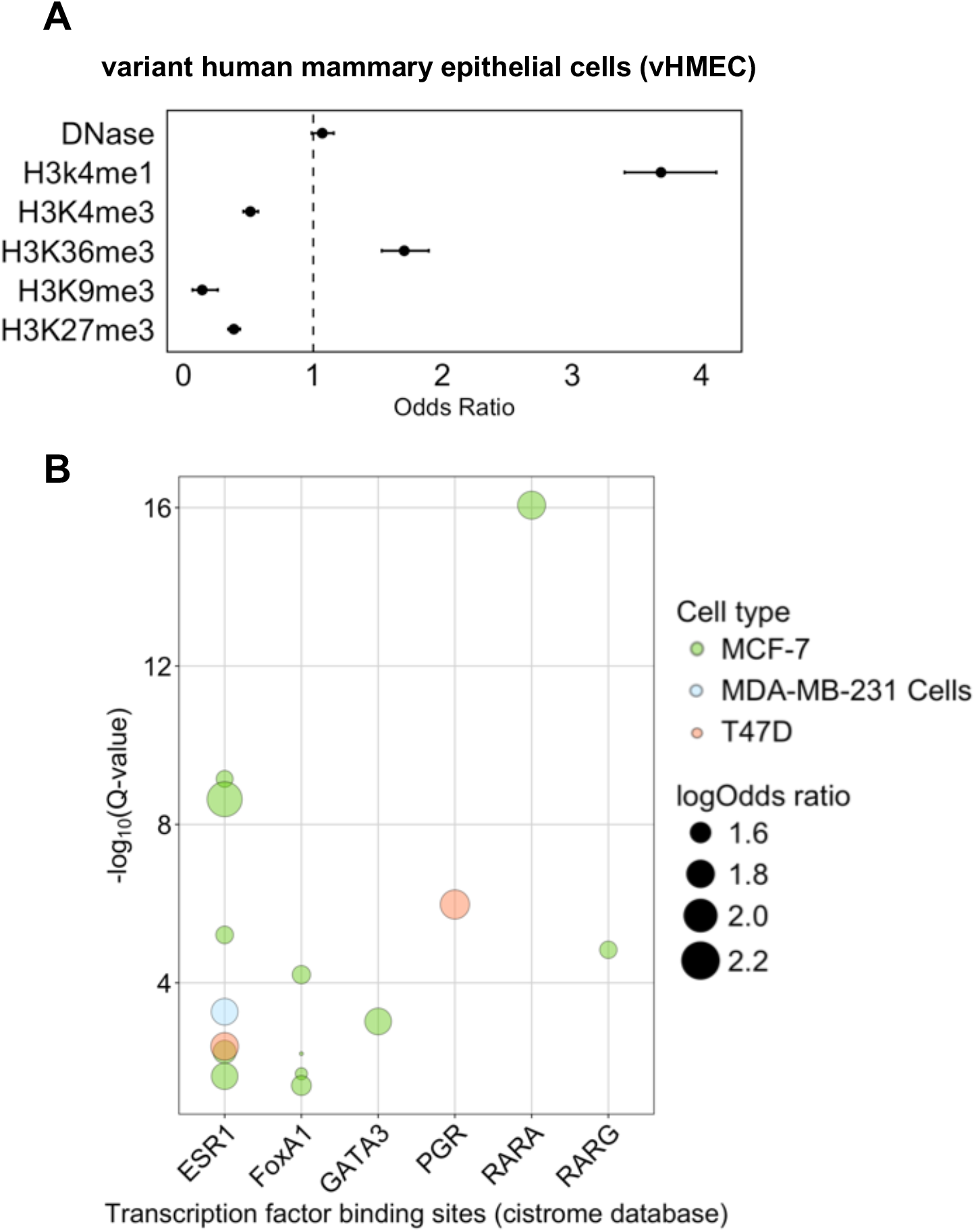
Normal breast tissue 5hmC is enriched at regulatory regions in transformed and malignant breast cell lines. (**A**) Forest plot of Cochran-Mantel-Haenszel odds ratios and 95% confidence intervals (95% CI) from enrichment tests of high 5hmC CpGs among indicated genomic features, against 450K background set, in variants human mammary epithelial cells (vHMEC). Numerical values for odds ratios and *P*-values are provided in the adjacent table. (**B**) Vertical axis provides -Log10 FDR Q-value from enrichment tests (Fisher’s exact test) of high 5hmC CpGs, compared to 450K background set, among transcription factor binding sites (TFBSs, profiled as part of the Cistrome project) indicated on horizontal axis. Colors relate to cell lines profiled by chromatin immunoprecipitation sequencing (ChIP-seq) experiments for the indicated TFBSs. Bubble size relates to effect size (log odds ratio) from enrichment tests.

To explore potential enrichment of 5hmC among regulatory regions in breast cancer cell lines, we utilized Locus Overlap Analysis (LOLA) software to test genomic coordinates of the high 5hmC CpGs for enrichment among known TFBS and regulatory regions from publicly available data collections. Enrichment of 5hmC among fifteen TFBS across three breast cancer cell lines was observed at FDR <0.05 (**Figure 5B, Supplementary Data 9**) from the cistrome collection^36^. Strikingly, binding sites for ESR1 (estrogen receptor 1) were enriched for 5hmC in MCF7, T47D, MDA-MB-231 cells (Figure 5B). Binding sites for other transcription factors involved in regulation of breast specific regulatory programs was also observed, including PGR (progesterone receptor), FOXA1, and GATA3 (**Figure 5B**). The LOLA algorithm was also used to query the enrichment of 5hmC among TFBS from the ENCODE project collection. These analyses revealed enrichment for 5hmC among c-Fos binding sites in ER-Src transformed normal-like breast cell line MCF10A, as well as GATA3 binding sites in MCF-7 cells (**Supplementary Data 9**). Finally, enrichment for 5hmC was observed among H3K4me1-marked regions in T47D cells from the cistrome-epigenome collection, suggesting enrichment among active and poised enhancers (**Supplementary Data 9**).

### Normal breast tissue 5hmC is enriched among CpG loci differentially methylated between normal and invasive breast tissue

Numerous studies have identified DNA methylation differences between normal breast tissue and invasive disease. However, due to technical constraints, prior work has not distinguished 5hmC from 5mC when comparing breast tumor to normal breast tissue. We sought to investigate whether high 5hmC loci identified in normal breast tissue were more likely to be altered in breast tumors than other CpG loci. First, at the same 3876 high 5hmC CpG loci we identified in normal breast tissue, we compared DNA methylation between normal (*n*=95) and invasive breast tissue (*n*=753) from subjects in the TCGA project. Of the 3572 CpG loci available for testing, 1712 (47.9%) were differentially methylated between normal and invasive tissue in limma models adjusted for subject age and adjusted for multiple testing using Bonferroni correction (**Figure 6**). 1072 (62.6%) of the significantly differentially methylated loci had negative limma coefficients, suggesting 5hmC is more commonly lost from high 5hmC loci during breast cancer development, consistent with the observation that 5hmC is depleted in cancer tissues^23, 24^. Next, using randomly selected CpGs in the normal versus tumor comparison we created a null *P*-value distribution (see methods) and observed that the *P*-values for high 5hmC CpGs as a distribution were significantly lower (Kolmogorov-Smirnov test, *P*<0.05, **Table 2, Supplementary Figure 6A**). In addition, as extensive changes in DNA methylation have been observed to occur in ductal carcinoma *in situ* (DCIS)^37^, we repeated the procedure above using DNA methylation data from adjacent normal *(*n*=11*) and DCIS tissue samples (*n*=28) obtained from the New Hampshire Mammography Network (NHMN). In the comparison of DCIS to normal DNA methylation analysis, high 5hmC CpG sites identified in normal breast again had a distribution of *P*-values significantly lower than randomly selected CpGs (*P*=0.017, **Table 2, Supplementary Figure 6B**). Together these results suggest 5hmC dysregulation may contribute to breast cancer development, and an appreciable fraction of the changes may occur early in the carcinogenic process.

**Figure 6.**
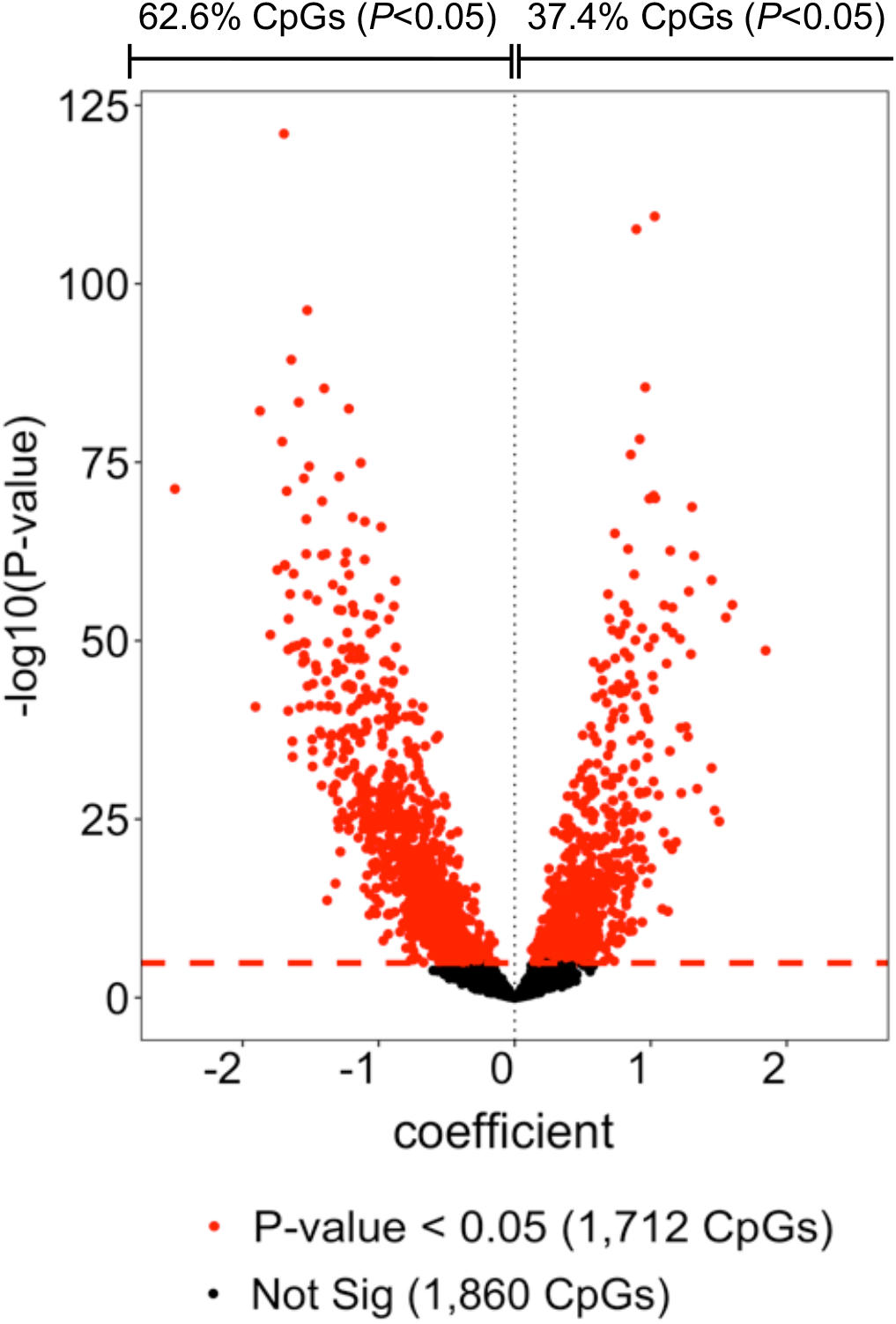
Differential methylation analysis of tumor and normal breast tissue at high 5hmc CpG loci. Volcano plot shows results from 3572 age-adjusted multivariable linear (limma) models of differential methylation across normal (*n* = 95) and tumor tissue (*n* = 753) for each of the high 5hmC CpGs. Red points demonstrated statistically significant differential methylation status across normal and tumor tissues after correction for multiple testing. Dashed red line indicates Bonferroni significance threshold for 3572 tests (*P* = 1.3E-05).

**Table 2.** Statistical comparison of *P*-value distributions obtained in differential methylation analyses between tumor/DCIS and adjacent normal tissues

| Data-set | Comparison | Kolmogorov-Smirnov test |  |
| --- | --- | --- | --- |
|  |  | *D*-statistic | *P*-value |
| TCGA | Normal vs tumor | 0.032 | 0.046 |
| NHMN | Normal vs DCIS | 0.036 | 0.017 |

Two-sided Kolmogorov-Smirnov (KS) tests were used to compare *P*-value distributions from a randomly generated CpG set (see Methods) with those obtained from differential methylation analyses in the indicated data sets. TCGA, The Cancer Genome Atlas; NHMN, New Hampshire Mammography Network.

## Discussion

While the distribution and functions of 5mC in various cellular contexts has been studied at depth, much less is known about the importance of other cytosine modifications. Traditional approaches utilizing DNA BS treatment to distinguish between cytosine and 5mC have been limited, as they are unable to disambiguate 5mC from its oxidized forms. Here, we performed tandem BS and oxBS treatment of DNA from normal breast tissue coupled with 450K arrays for DNA methylation to estimate relative 5mC and 5hmC proportions and construct a genome-wide map of 5hmC in breast tissue. 5hmC was generally depleted compared with 5mC as expected, but specific loci with recurrent 5hmC across individual samples were identified. Regions containing abundant 5hmC were enriched among enhancers and transcriptionally active chromatin regions and depleted in transcriptionally repressed areas of the genome. Furthermore, we provide evidence suggesting deregulation of 5hmC at these loci may contribute to breast carcinogenesis. Collectively, our findings reveal 5hmC is present and uniquely distributed in normal breast tissue and may hold gene regulatory functions protecting against carcinogenesis.

Previous studies have observed a lack of obvious correlation between levels of 5mC and forms of oxidized cytosine in individual tissues^4, 38^. Such findings suggest the abundance of oxidized cytosine may be subject to a higher order regulation in a tissue-specific context. Indeed, active DNA demethylation controls various cellular processes such as somatic cell reprogramming^39, 40^, chromatin accessibility^11, 13, 41^, as well as other processes involved in cell fate specification^42, 43, 44^. Enrichment of 5hmC among enhancers^6, 9, 32^, DNase hypersensitivity sites^9^, and gene bodies^9, 32, 33, 45^ observed in multiple cell types supports the proposal that 5hmC is a functional epigenetic mark involved in positive regulation of gene expression. Consistent with these findings, we found 5hmC enrichment among introns, enhancers, and regions with transcriptionally active chromatin. Representation of multiple transcriptional co(activators) and chromatin regulators, some of which hold established functions in breast tissue, among the genes with greatest 5hmC abundance suggests involvement of 5hmC in breast tissue-specific gene expression control. In contrast to prior studies of other tissue types, we did not observe enrichment for 5hmC among breast-specific DNase hypersensitivity sites. Gene set enrichment analyses of those genes containing the most abundant 5hmC suggested potential function for 5hmC in regulation of immune cell activity, TGF-beta regulation, and cell motility. Collectively, these findings reveal a unique distribution of 5hmC in normal breast tissue that warrants further study.

Although decrease of total 5hmC content has been observed invariably over a wide range of cancers^24, 46, 47^, the contribution of locus-specific changes in 5hmC levels to cancer pathogenesis is not well understood. Emerging evidence suggests acquisition of 5hmC among enhancer regions and disease-relevant oncogenes promotes growth and survival of tumor cells, as has been observed in pancreatic ductal adenocarcinoma, glioblastoma, and breast tumor initiating cells^33, 48, 49, 50^. Similarly, our findings that normal breast tissue 5hmC is enriched among enhancer regions in vHMEC cells, suggests maintenance of enhancer-specific 5hmC may be required for maintenance of their premalignant characteristics. In addition, our observation that normal breast 5hmC is enriched among estrogen receptor alpha (*ESR1*) binding sites in multiple breast cancer cell lines suggests 5hmC may contribute to disease-relevant processes. In addition to *ESR1*, itself a critical determinant of estrogen therapy treatment response in ER^+^ breast cancer^51^, our findings that 5hmC is enriched among binding sites for *FOXA1, GATA3*, and the *RAR* family transcription factors, all known to function cooperatively with *ESR1* in normal and malignant breast tissue^52, 53, 54^, further supports the notion that locus-specific maintenance of 5hmC contributes to breast cancer pathogenesis. Whether CpG loci within these regulatory regions are subject to locus-specific increases in 5hmC or are protected from 5hmC loss during carcinogenesis will require more detailed interrogation of 5hmC at these loci across normal and malignant tissues. Additionally, our observation that the high 5hmC CpGs identified in this cohort are enriched among differentially methylated loci between normal and malignant tissue supports the idea that alterations in 5hmC contribute to breast cancer pathogenesis. Future work is needed to examine the directionality of this phenomenon at the level of individual genes. Total genomic 5hmC in breast tumors has also been observed as a poor prognostic indicator^25^, suggesting that 5hmC abundance may also contribute to tumor progression. Studies profiling 5hmC at base-resolution in tumors with available follow up and outcome data are required to further explore this relationship.

While we are encouraged by the consistency of our findings with previously published work, our measurements of 5hmC are aggregates across breast tissue cell types, which could introduce noise into our data. Though we addressed this issue using a reference-free cell type deconvolution method, we recognize that larger studies will provide more precise estimates of cell type proportions and their potential variation across subjects. Additionally, although the array-based approach we utilized is an order of magnitude less costly than whole-genome bisulfite sequencing, we expect such an approach would provide additional insights to the distribution and functional implications of 5hmC in breast tissue in future studies.

In summary, we analyzed data from tandem processed BS and oxBS-treated methylation arrays to produce a genome-wide map of 5hmC and 5mC in normal breast tissue. Identification of multiple transcriptional (co)-activators and chromatin modifying genes containing elevated levels of 5hmC, including several breast-specific TSGs, suggests 5hmC may be involved in regulation of key transcriptional programs in normal breast tissue. Consistent with previous findings, 5hmC was highly enriched among regulatory regions associated with transcriptionally active chromatin, while depleted among regions associated with gene repression. Additionally, we provide evidence that dysregulation of 5hmC in breast tissue may contribute to carcinogenesis. These findings extend our understanding of epigenetic regulation in normal breast tissue and provide a reference that can be used in the study of functions of 5hmC in diseased breast tissue.

## Methods

### Study population

Fresh-frozen disease-free breast tissue was obtained from 18 subjects sourced from the National Disease Research Interchange (NDRI). All subjects provided written informed consent at the time of surgery for use of their tissue specimens in research as approved by the committee for protection of human subjects (IRB). Subject demographics were available for study and all tissues accessed were from deceased subjects. This work was performed in accordance with the ethical principles in the Declaration of Helsinki.

### DNA extraction, conversion, and methylation profiling

DNA extraction was performed with the QIAmp DNeasy tissue kit (Qiagen) according to the manufacturer’s instructions. DNA conversion and methylation profiling have been described previously. Briefly, quantity and quality of breast tissue DNA was determined with the Qubit 3.0 fluorometer (Life Technologies). Tandem bisulfite and oxidative bisulfite conversion was performed using the TrueMethyl^®^ kit v.1.1 (Cambridge Epigenetix) protocol optimized for Infinium HumanMethylation450 arrays (450K, Illumina, Inc., San Diego, CA), with an input of 4ug per sample. Genomic DNA was then sheared to ~10kb fragments using g-TUBE (Covaris) and purified with the Gene-JET PCR Purification kit (Thermo Scientific). oxBS conversion was then performed with 1.4ug of sheared DNA according to the TrueMethyl protocol, and 1.05ug for bisulfite conversion using manufacturer recommended mass and volume. ssDNA was recovered and quantified with Qubit and processed on Illumina 450K methylation arrays at the UCSF genomics core facility.

### RNA extraction and gene expression

RNA extraction was performed using the RNeasy Mini tissue kit (Qiagen) according to the manufacturer’s instructions. Qubit 3.0 was used to determine RNA quality and quantity prior to expression profiling. Absolute expression quantification was performed using the nCounter Analysis System (NanoString Technologies). Transcripts selection was performed in conjunction with previously published work^33^, and limited to genes with known functions in breast tissue and an appreciable abundance of 5hmC, as well as being amenable to probe design for the nCounter assay. Epigenetic enzymes *TET1, TET2, TET3, DNMT3A* (transcript variants 2, 3, 4), *DNMT3B*, and *DNMT1* were also selected for gene expression profiling. Platform associated variation was normalized using the nSolver Analysis software (NanoString, V2.6). Expression of candidate transcripts of interest was normalized to that of housekeeping genes *PUM1, BUSB, TBP, ACTB*, and *SDHA*. Normalized gene expression data from this experiment are available in **Supplementary Data 6**.

### Data processing and statistical analysis

#### 5-(hydroxy)methylcytosine estimation and quality control

All data analysis was conducted in R version 3.3.1. Data processing of raw signals from BS and oxBS-treated samples has been previously described. Briefly, normalization and background correction were performed using the *FunNorm* procedure available in the R/Bioconductor package *minfi* (version 1.10.2). CpG sites located on sex chromosome or those previously identified as cross-reactive or containing SNPs were removed prior to analysis. After quality control, 387,617 CpGs were left for analysis. To perform estimation of 5mC and 5hmC proportions in each sample, we applied the recently developed OxyBS algorithm, which uses a maximum likelihood-based method and more appropriate statistical constraints than previous methods to accurately predict methylated and unmethylated proportions. Code used to perform the estimation is available in R-package ‘OxyBS’^29^. To identify CpG sites with greatest potential to be functionally relevant, we selected the 3876 CpGs among the top 1% median 5hmC value across all samples. These CpGs are referred to as the ‘high 5hmC CpGs’ in the results section.

#### Enrichment analyses of high 5hmC CpG loci

To test genomic feature enrichment, the *GenomicRanges* R/Bioconductor package was used to constructed contingency tables describing the overlap between genomic coordinates of high 5hmC CpGs and specific genomic features of interest (for example, post-translationally modified histones), stratified by CpG island context. This approach resulted in four contingency tables per enrichment test, each denoting the overlap between high 5hmC CpGs and the genomic feature within that specific CpG island context. We then applied Cochran-Mantel-Haenszel tests to obtain odds ratios (ORs) and *P*-values. For all enrichment tests, CpG sites assayed on the Infinium 450K array were used as background. Genomic regions (introns, exons, promoters, intergenic) were defined using the UCSC_hg19_refGene file from the UCSC Genome Browser. Where CpG loci were associated with more than one genomic regions, the following precedence was applied: promoter>exon>intron. R/Bioconductor packages *IlluminaHumanMethylation450kmanifest*, version 0.4.0, and IlluminaHumanMethylation450kanno.ilmn12.hg19, version 0.2.1 to obtain relevant array annotation data, including CpG island context.

#### Preprocessing and quality control of TCGA and GSE66313 data-sets

DNA methylation data from the TCGA project and NHMN (GSE66313) were subjected to the preprocessing and quality control pipeline described in Salas *et al*^55^. Briefly, raw intensity data files (IDATs) obtained using the 450K microarray platform were imported and processed using the *RnBeads* R package^56^. Methylation β-values for individual CpG loci were calculated as the ratio of methylated probe intensity divided by the total signal from the methylated and unmethylated signal intensities, plus an offset of 100^57^. Background correction was performed using methylumi-noob^58^ and normalized using a functional normalization procedure^59^. Probes designed against CpG loci on sex chromosomes, non CpG loci, or previously documented as polymorphic or crossreactive were excluded from subsequent analyses^60^. Finally, the Greedycut hierarchical algorithm^56^ was applied to the remaining data to identify and remove unreliable samples/probes. Briefly, Greedycut iteratively produces a matrix of retained and removed measurements each time the algorithm is applied to data where probes/samples are iteratively added/removed. Probes/samples with the highest fraction of unreliable measurements are removed from further analysis.

#### Analysis of CpG specific associations

Differential methylation status between normal-adjacent and tumor/DCIS tissue at individual CpG-loci in the TCGA and NHMN data sets was determined by using multivariable linear models for microarray data (limma)^61^ to model logit-transformed methylation β-values (*M*-values). Models were adjusted for subject age, and Bonferroni correction was used to adjust for multiple testing. To determine if the proportion of high 5hmC CpG loci demonstrating significant differential methylation between normal and invasive breast tissue was greater than would be expected due to chance, we first took 1000 samples of randomly selected CpG loci of identical size (*n*=3572) and CpG island context to the high 5hmC CpG loci. After ordering each randomly selected CpG set according to *P*-value magnitude, a single *P*-value distribution representative of a ‘null’ distribution was generated by averaging *P*-values across the 1000 sets. *P*-value distributions for the high-5hmC loci and the set representative of the ‘null’ distribution was then compared using the Kolmogorov-Smirnov test (**Supplementary Figure 6**).

#### Cell-mixture deconvolution

Fluctuations in cell-type proportions between samples is a well-documented potential confounder in studies of DNA methylation^62, 63^. Reference-free cell mixture deconvolution methods have been recently developed and widely used^64^, to infer putative cell type proportions in studies of heterogeneous tissues where tissue-specific reference DNA methylomes do not exist. We used the RefFreeEWAS algorithm, implemented using R-package *RefFreeEWAS*^31^ to estimate putative cell-types and their cellular proportions in each sample. A variant of non-negative matrix factorization, the RefFreeEWAS algorithm attempts to identify the major axes of cellular variation in DNA methylation data and deconvolute these to methylomes of the individual cell types. Using methylation proportions from BS-treated DNA, we selected the 10,000 most variable CpG loci to determine the optimal number of cell-types (K) that explain the methylation data across all 18 samples. K=2 was identified as the optimal number of putative cell-types. Finally, the full set of 387,617 passing the quality control procedures described above, were used to obtain sample-specific estimates of the proportions for each of the two putative cell-types.

## Data Availability

Raw (IDAT files) and normalized DNA methylation data (450K platform) used to estimate 5hmC and 5mC propotions from the 18 normal breast tissue samples are available under accession GSE73895 in the Gene Expression Omnibus (GEO) (http://www.ncbi.nlm.nih.gov/geo/). Raw DNA methylation data for DCIS and adjacent-normal tissue samples from the NHMN is also available in GEO (GSE66313). Level 1 TCGA breast (BRCA) intensity files derived from the 450K platform were downloaded from the TCGA data portal and are currently available through the National Cancer Institute (NCI) Genomic Data Commons (GDC) data portal (https://portal.gdc.cancer.gov/). Coordinates of DNase hypersensitivity sites and histone modfications (derived from ChIP-seq experiments) in breast myoepiethial cells and HMECs were downloaded from the NIH Roadmap Epigenomics Project^65^ and are also available in GEO. Specific GEO accession numbers from each Roadmap Epigenomics experiment can be found at https://www.ncbi.nlm.nih.gov/geo/roadmap/epigenomics/. All other data are available within the article or Supporting Information.

## Code Availability

R code used for all analyses is available in the ‘Normal-Breast-5hmC’ repository on GitHub (https://github.com/Christensen-Lab-Dartmouth).

## Financial Information

This work was supported by the National Institutes of Health grants R01DE022772 and R01CA216265 [to B.C.C.], R01MH094609 [to E.A.H], and the Center for Molecular Epidemiology COBRE program funded by P20GM104416.

## Author Contributions

O.M.W. conceived and designed the approach, performed statistical analyses, interpreted the results, wrote and revised the manuscript. K.C.J. conceived and designed the approach, performed statistical analyses, interpreted the results, wrote and revised the manuscript. E.A.H. conceived and designed the approach, generated statistical models, performed statistical analyses, interpreted the results, and revised the manuscript. J.E.K. carried out laboratory experiments and revised the manuscript. C.J.M. conceived and designed the approach and revised the manuscript. B.C.C. conceived and designed the approach, interpreted the results, wrote and revised the manuscript. All authors have read and approved the final manuscript.

## References

1. Jones PA. Functions of DNA methylation: islands, start sites, gene bodies and beyond. Nat Rev Genet 13, 484–492 (2012).

2. Kriaucionis S, Heintz N. The nuclear DNA base 5-hydroxymethylcytosine is present in Purkinje neurons and the brain. Science 324, 929–930 (2009).

3. Tahiliani M, et al. Conversion of 5-methylcytosine to 5-hydroxymethylcytosine in mammalian DNA by MLL partner TET1. Science 324, 930–935 (2009).

4. Ito S, et al. Tet proteins can convert 5-methylcytosine to 5-formylcytosine and 5-carboxylcytosine. Science 333, 1300–1303 (2011).

5. Wu X, Zhang Y. TET-mediated active DNA demethylation: mechanism, function and beyond. Nat Rev Genet 18, 517–534 (2017).

6. Sun Z, et al. A sensitive approach to map genome-wide 5-hydroxymethylcytosine and 5-formylcytosine at single-base resolution. Mol Cell 57, 750–761 (2015).

7. Pastor WA, et al. Genome-wide mapping of 5-hydroxymethylcytosine in embryonic stem cells. Nature 473, 394–397 (2011).

8. Ficz G, et al. Dynamic regulation of 5-hydroxymethylcytosine in mouse ES cells and during differentiation. Nature 473, 398–402 (2011).

9. Yu M, et al. Base-resolution analysis of 5-hydroxymethylcytosine in the mammalian genome. Cell 149, 1368–1380 (2012).

10. Wen L, et al. Whole-genome analysis of 5-hydroxymethylcytosine and 5-methylcytosine at base resolution in the human brain. Genome Biol 15, R49 (2014).

11. Iurlaro M, et al. A screen for hydroxymethylcytosine and formylcytosine binding proteins suggests functions in transcription and chromatin regulation. Genome Biol 14, R119 (2013).

12. Spruijt CG, et al. Dynamic readers for 5-(hydroxy)methylcytosine and its oxidized derivatives. Cell 152, 1146–1159 (2013).

13. Mellen M, Ayata P, Dewell S, Kriaucionis S, Heintz N. MeCP2 binds to 5hmC enriched within active genes and accessible chromatin in the nervous system. Cell 151, 1417–1430 (2012).

14. Zhong J, et al. TET1 modulates H4K16 acetylation by controlling auto-acetylation of hMOF to affect gene regulation and DNA repair function. Nucleic Acids Res 45, 672–684 (2017).

15. Kafer GR, et al. 5-Hydroxymethylcytosine Marks Sites of DNA Damage and Promotes Genome Stability. Cell Rep 14, 1283–1292 (2016).

16. Nestor CE, et al. Tissue type is a major modifier of the 5-hydroxymethylcytosine content of human genes. Genome Res 22, 467–477 (2012).

17. Jin S-G, Wu X, Li AX, Pfeifer GP. Genomic mapping of 5-hydroxymethylcytosine in the human brain. Nucleic Acids Res 39, 5015–5024 (2011).

18. Guo JU, Su Y, Zhong C, Ming G-l, Song H. Hydroxylation of 5-methylcytosine by TET1 promotes active DNA demethylation in the adult brain. Cell 145, 423–434 (2011).

19. Song C-X, et al. Selective chemical labeling reveals the genome-wide distribution of 5-hydroxymethylcytosine. Nat Biotechnol 29, 68–72 (2011).

20. Plachot C, Lelievre SA. DNA methylation control of tissue polarity and cellular differentiation in the mammary epithelium. Exp Cell Res 298, 122–132 (2004).

21. Fleischer T, et al. Genome-wide DNA methylation profiles in progression to in situ and invasive carcinoma of the breast with impact on gene transcription and prognosis. Genome Biol 15, 435 (2014).

22. Johnson KC, et al. DNA methylation in ductal carcinoma in situ related with future development of invasive breast cancer. Clin Epigenetics 7, 75 (2015).

23. Jin S-G, et al. 5-Hydroxymethylcytosine is strongly depleted in human cancers but its levels do not correlate with IDH1 mutations. Cancer Res 71, 7360–7365 (2011).

24. Kudo Y, et al. Loss of 5-hydroxymethylcytosine is accompanied with malignant cellular transformation. Cancer Sci 103, 670–676 (2012).

25. Tsai K-W, et al. Reduction of global 5-hydroxymethylcytosine is a poor prognostic factor in breast cancer patients, especially for an ER/PR-negative subtype. Breast Cancer Res Treat 153, 219–234 (2015).

26. Song SJ, et al. MicroRNA-antagonism regulates breast cancer stemness and metastasis via TET-family-dependent chromatin remodeling. Cell 154, 311–324 (2013).

27. Thienpont B, et al. Tumour hypoxia causes DNA hypermethylation by reducing TET activity. Nature 537, 63–68 (2016).

28. Li L, et al. Epigenetic inactivation of the CpG demethylase TET1 as a DNA methylation feedback loop in human cancers. Sci Rep 6, 26591 (2016).

29. Houseman EA, Johnson KC, Christensen BC. OxyBS: estimation of 5-methylcytosine and 5-hydroxymethylcytosine from tandem-treated oxidative bisulfite and bisulfite DNA. Bioinformatics 32, 2505–2507 (2016).

30. Jaffe AE, Irizarry RA. Accounting for cellular heterogeneity is critical in epigenome-wide association studies. Genome Biol 15, R31 (2014).

31. Houseman EA, Molitor J, Marsit CJ. Reference-free cell mixture adjustments in analysis of DNA methylation data. Bioinformatics 30, 1431–1439 (2014).

32. Wu H, et al. Genome-wide analysis of 5-hydroxymethylcytosine distribution reveals its dual function in transcriptional regulation in mouse embryonic stem cells. Genes Dev 25, 679–684 (2011).

33. Johnson KC, Houseman EA, King JE, von Herrmann KM, Fadul CE, Christensen BC. 5-Hydroxymethylcytosine localizes to enhancer elements and is associated with survival in glioblastoma patients. Nat Commun 7, 13177 (2016).

34. Stroud H, Feng S, Morey Kinney S, Pradhan S, Jacobsen SE. 5-Hydroxymethylcytosine is associated with enhancers and gene bodies in human embryonic stem cells. Genome Biol 12, R54 (2011).

35. Ivanov M, et al. Ontogeny, distribution and potential roles of 5-hydroxymethylcytosine in human liver function. Genome Biol 14, R83 (2013).

36. Liu T, et al. Cistrome: an integrative platform for transcriptional regulation studies. Genome Biol 12, R83 (2011).

37. Fleischer T, et al. Genome-wide DNA methylation profiles in progression to in situ and invasive carcinoma of the breast with impact on gene transcription and prognosis. Genome Biol 15, 435 (2014).

38. Bachman M, et al. 5-Formylcytosine can be a stable DNA modification in mammals. Nat Chem Biol 11, 555–557 (2015).

39. Doege CA, et al. Early-stage epigenetic modification during somatic cell reprogramming by Parp1 and Tet2. Nature 488, 652–655 (2012).

40. Hu X, et al. Tet and TDG mediate DNA demethylation essential for mesenchymal-to-epithelial transition in somatic cell reprogramming. Cell Stem Cell 14, 512–522 (2014).

41. Lio C-W, Zhang J, Gonzalez-Avalos E, Hogan PG, Chang X, Rao A. Tet2 and Tet3 cooperate with B-lineage transcription factors to regulate DNA modification and chromatin accessibility. Elife 5, (2016).

42. Wheldon LM, et al. Transient accumulation of 5-carboxylcytosine indicates involvement of active demethylation in lineage specification of neural stem cells. Cell Rep 7, 1353–1361 (2014).

43. Yue X, et al. Control of Foxp3 stability through modulation of TET activity. J Exp Med 213, 377–397 (2016).

44. Orlanski S, et al. Tissue-specific DNA demethylation is required for proper B-cell differentiation and function. Proc Natl Acad Sci U S A 113, 5018–5023 (2016).

45. Xu Y, et al. Genome-wide regulation of 5hmC, 5mC, and gene expression by Tet1 hydroxylase in mouse embryonic stem cells. Mol Cell 42, 451–464 (2011).

46. Haffner MC, et al. Global 5-hydroxymethylcytosine content is significantly reduced in tissue stem/progenitor cell compartments and in human cancers. Oncotarget 2, 627–637 (2011).

47. Lian CG, et al. Loss of 5-hydroxymethylcytosine is an epigenetic hallmark of melanoma. Cell 150, 1135–1146 (2012).

48. Bhattacharyya S, et al. Altered hydroxymethylation is seen at regulatory regions in pancreatic cancer and regulates oncogenic pathways. Genome Res 27, 1830–1842 (2017).

49. Takai H, et al. 5-Hydroxymethylcytosine plays a critical role in glioblastomagenesis by recruiting the CHTOP-methylosome complex. Cell Rep 9, 48–60 (2014).

50. Wu M-Z, et al. Hypoxia Drives Breast Tumor Malignancy through a TET-TNFalpha-p38-MAPK Signaling Axis. Cancer Res 75, 3912–3924 (2015).

51. Ross-Innes CS, et al. Differential oestrogen receptor binding is associated with clinical outcome in breast cancer. Nature 481, 389–393 (2012).

52. Hurtado A, Holmes KA, Ross-Innes CS, Schmidt D, Carroll JS. FOXA1 is a key determinant of estrogen receptor function and endocrine response. Nat Genet 43, 27–33 (2011).

53. Theodorou V, Stark R, Menon S, Carroll JS. GATA3 acts upstream of FOXA1 in mediating ESR1 binding by shaping enhancer accessibility. Genome Res 23, 12–22 (2013).

54. Hua S, Kittler R, White KP. Genomic antagonism between retinoic acid and estrogen signaling in breast cancer. Cell 137, 1259–1271 (2009).

55. Salas LA, Johnson KC, Koestler DC, O'Sullivan DE, Christensen BC. Integrative epigenetic and genetic pan-cancer somatic alteration portraits. Epigenetics 12, 561–574 (2017).

56. Assenov Y, Muller F, Lutsik P, Walter J, Lengauer T, Bock C. Comprehensive analysis of DNA methylation data with RnBeads. Nat Methods 11, 1138–1140 (2014).

57. Du P, et al. Comparison of Beta-value and M-value methods for quantifying methylation levels by microarray analysis. BMC Bioinformatics 11, 587 (2010).

58. Triche TJ, Jr., Weisenberger DJ, Van Den Berg D, Laird PW, Siegmund KD. Low-level processing of Illumina Infinium DNA Methylation BeadArrays. Nucleic Acids Res 41, e90 (2013).

59. Fortin JP, et al. Functional normalization of 450k methylation array data improves replication in large cancer studies. Genome Biol 15, 503 (2014).

60. Chen YA, et al. Discovery of cross-reactive probes and polymorphic CpGs in the Illumina Infinium HumanMethylation450 microarray. Epigenetics 8, 203–209 (2013).

61. Ritchie ME, et al. limma powers differential expression analyses for RNA-sequencing and microarray studies. Nucleic Acids Res 43, e47 (2015).

62. Houseman EA, et al. DNA methylation arrays as surrogate measures of cell mixture distribution. BMC Bioinformatics 13, 86 (2012).

63. Houseman EA, Kelsey KT, Wiencke JK, Marsit CJ. Cell-composition effects in the analysis of DNA methylation array data: a mathematical perspective. BMC Bioinformatics 16, 95 (2015).

64. Titus AJ, Gallimore RM, Salas LA, Christensen BC. Cell-type deconvolution from DNA methylation: a review of recent applications.Hum Mol Genet 26, R216–R224 (2017).

65. Bernstein BE, et al. The NIH Roadmap Epigenomics Mapping Consortium. Nat Biotechnol 28, 1045–1048 (2010).

